# HIV-1 Vif gained breadth in APOBEC3G specificity after cross-species transmission of its precursors

**DOI:** 10.1101/2021.08.23.457421

**Authors:** Nicholas M. Chesarino, Michael Emerman

**Author notes:** Corresponding author. Michael Emerman, 1100 Fairview Avenue N., C2-023, Seattle WA, USA 98109.

## Abstract

APOBEC3G (A3G) is a host-encoded cytidine deaminase that potently restricts retroviruses, such as HIV-1, and depends on its ability to package into virions. As a consequence of this, HIV-1 protein Vif has evolved to antagonize human A3G by targeting it for ubiquitination and subsequent degradation. There is an ancient arms-race between Vif and A3G highlighted by amino acids 128 and 130 in A3G that have evolved under positive selection due to Vif-mediated selective pressure in Old World primates. Nonetheless, not all possible amino acid combinations at these sites have been sampled by nature and it is not clear the evolutionary potential of species to resist Vif antagonism. To explore the evolutionary space of positively selected sites in the Vif-binding region of A3G, we designed a combinatorial mutagenesis screen to introduce all 20 amino acids at sites 128 and 130. Our screen uncovered mutants of A3G with several interesting phenotypes, including loss of stability and resistance of Vif antagonism. However, HIV-1 Vif exhibited remarkable flexibility in antagonizing A3G 128 and 130 mutants, which significantly reduces viable Vif resistance strategies for hominid primates. Importantly, we find that broadened Vif specificity was conferred through Loop 5 adaptations that were required for cross-species adaptation from Old World monkey A3G to hominid A3G. Our evidence suggests that Vif adaptation to novel A3G interfaces during cross-species transmission may train Vif towards broadened specificity that can further facilitate cross-species transmissions and raise the barrier to host resistance.

## 1. Introduction

Many of the emergent viral epidemics in the human population are the result of cross-species, or zoonotic, transmission from animals. One of the most notable of these is human immunodeficiency virus-1 (HIV-1), which became pandemic following successful cross-species transmission of simian immunodeficiency virus (SIV) infecting chimpanzees (SIVcpz). SIVcpz itself resulted from a series of recombination and cross-species transmission events involving an SIV infecting red-capped mangabeys (SIVrcm) and an SIV infecting the guenons [1,2]. In addition, a region at the 5’ end of the SIVcpz/HIV-1 lineage has not been accounted for [3,4]. Importantly, recombination between lentiviral lineages was likely not sufficient to establish productive infection in hominid primates, as further molecular adaptations took place to antagonize and evade host antiviral factors in chimpanzees [3,5].

Host antiviral factors with cell-intrinsic antiviral activity, often called restriction factors, have the capacity to block replication of viruses at a variety of stages of the viral lifecycle, and act as the first line of defense against these intracellular parasites [6,7]. Importantly, these antiviral proteins establish a network of barriers that effectively block cross-species transmission of maladapted viruses, and establishment of a virus into a new host requires overcoming restriction factors by way of evasion or direct antagonism [8,9]. Thus, viral genes must adapt binding interfaces to block or enable interaction with divergent restriction factors while maintaining canonical functional roles.

As a result of a longstanding genetic conflict with retroviruses and other endogenous and exogenous pathogens over millions of years, primate *APOBEC3* genes are rapidly evolving under positive selection [10–14]. Positive selection of APOBEC3G (here, abbreviated as A3G), the most potent APOBEC3 against primate lentiviruses, has been shown to influence its interaction with lentiviral Vif [15–17]. Two of these sites in A3G, amino acids at positions 128 and 130, directly govern species-specific recognition by most Vif proteins [15–23]. Importantly, the A3G D128 residue found in all hominids confers resistance to many Old World monkey SIV Vif proteins, which largely recognize K128 present in nearly all Old World monkey A3G proteins [3,15,19–22]. Therefore, successful zoonosis of SIV from Old World monkeys to hominid primates involved adaptation of these SIV Vif proteins to the hominid (D128) version of A3G [3,5,24].

We recently described the primary amino acid substitution in SIVrcm Vif, Y86H, that conferred adaptation towards the D128 residue found exclusively in hominid A3G [5]. This change was required for SIVrcm Vif, the precursor to SIVcpz Vif, to fully antagonize chimpanzee A3G, and no further changes were needed for this mutant to antagonize human A3G. Moreover, we found that this SIVrcm Vif Y86H mutant remained fully active against rcmA3G, indicating that adaptation to hominid A3G did not come at a cost to host A3G antagonism [5]. Thus, the Y86H mutation in SIVrcm Vif broadened specificity of Vif towards the simultaneous antagonism of rcmA3G and hominid A3G, suggesting that this adaptive mutation in Vif may also confer greater tolerance of prospective changes at 128 and 130. Here, we explore how broadened HIV-1 Vif specificity impacts prospective resistance strategies for human A3G.

In this study, we created libraries of amino acid mutations at positions 128 and 130 in A3G to explore the prospective capacity of human A3G to resist HIV-1 Vif antagonism. While several strategies for HIV-1 Vif resistance are possible, we find that HIV-1 Vif can maintain antagonism against many human A3G mutants, especially those that are most evolutionarily accessible. We determined that Vif evolved broadened substrate specificity by way of Loop 5 adaptations to hominid primate A3G during the cross-species adaptation of SIVrcm Vif to chimpanzee A3G. In addition, not all pathways towards resistance from Vif are possible as some amino acid mutations at 128/130 lose antiviral activity. Our work demonstrates HIV-1 Vif developed broadened specificity towards A3G as a consequence of previous host adaptations and suggests that hominid primates are at an evolutionary disadvantage in the host-virus arms race against Vif due to their fixation of amino acids not amenable to Vif resistance.

## 2. Results

### 2.1. Implementation of a combinatorial mutagenesis screen to evaluate A3G-Vif antagonism

Previous studies have shown that a single amino acid change in human A3G, D128K, is sufficient to resist HIV-1 Vif antagonism without impacting antiviral efficacy [19–21]. Similarly, single amino acid changes at either 128 or 130 in Old World monkey A3G are sufficient to resist SIV Vif antagonism [15,22,23,25]. While these residues govern A3G specificity with many lentiviral Vif proteins, the residues found in primate A3G at sites 128 and 130 represent only a small subset of all possible amino acids at this position. Amino acids lysine (K), aspartic acid (D) and glutamic acid (E) have been found in primate A3G at residue 128, while amino acids aspartic acid (D), histidine (H), asparagine (N) and alanine (A) are found at residue 130 (**Figure 1A**). Of these residues, seven total combinations have been sampled by primate evolution of A3G, and these combinations retain antiviral efficacy while dramatically influencing interaction with lentiviral Vif [15,22].

**Figure 1.**
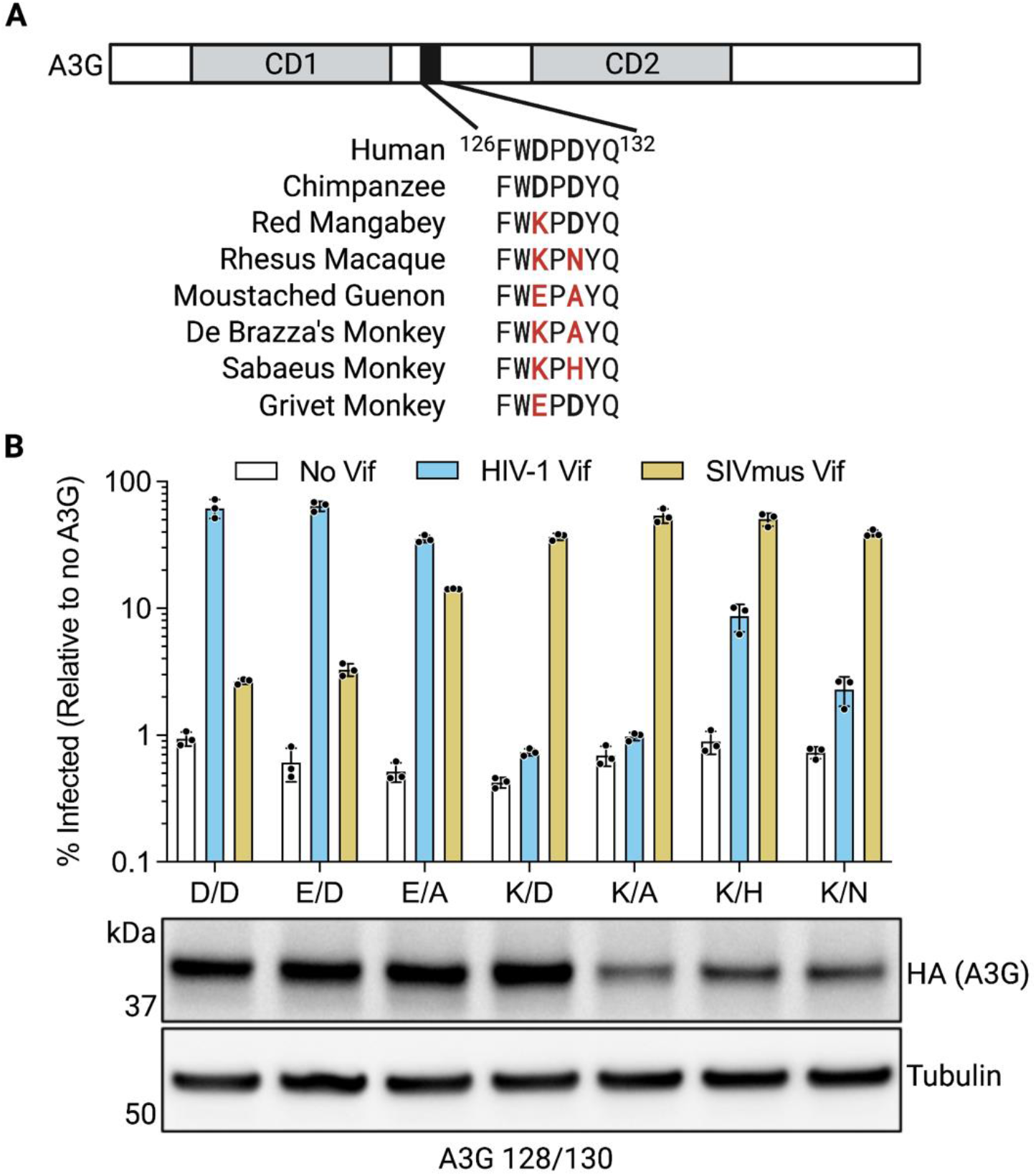
Extant mutations in human A3G 128 and 130 influence HIV-1 Vif susceptibility. **A**) Schematic of the primate A3G protein and regions of interest, including the first and second cytidine deaminase domains (CD1 and CD2, respectively) and the region between 126 and 132 that determine specificity for Vif (black box). Examples of primate A3G sequences represent all variants in this region found in nature (Compton *et al*., 2012; Compton and Emerman, 2013). Bold letters represent sites of positive selection, and red letters designate variations distinct from human A3G. **B**) Top: Single-cycle infectivity assays performed with HIV-1 without Vif (white), with Vif intact (blue), or with SIVmus Vif (gold) produced in the presence of indicated A3G 128/130 mutants. D/D denotes wild-type A3G at positions 128 and 130. Percent infectivity calculated relative to a no A3G control. Error bars represent the standard deviation of the mean of three biological replicates. Bottom: Immunoblot of cell lysate expressing wild-type A3G and the indicated A3G 128/130 mutant, performed using anti-HA to detect HA-tagged A3G, and anti-tubulin as a loading control.

To understand how the extant variation among Old World Monkeys at these positions might affect human A3G function and antagonism by HIV-1 Vif, we introduced these 128 and 130 variants into the human A3G background, transfected them into 293T cells, and assessed their ability to inhibit HIV-1 both in the presence and absence of Vif. D128E, D128E/D130A, and D128K mutants were expressed at levels consistent with wild-type, while expression of mutants D128K/D130N, D128K/D130H, and D128K/D130A was slightly reduced (**Figure 1B**). Despite differences in expression, all mutants display comparable antiviral efficacy in the absence of Vif (**Figure 1B**, infection reduced to ~1% compared to no A3G control). On the other hand, the ability of these A3G variants to be antagonized by HIV-1 Vif is strongly influenced by the identity of amino acid 128 – a negative residue (D, E) is potently antagonized by Vif, whereas K128 resists Vif (**Figure 1B**). Interestingly, the D128K/D130H mutant is weakly antagonized by HIV-1 Vif, demonstrating that residue 130 does influence human A3G susceptibility to Vif. Mutants not antagonized by HIV-1 Vif can still be recognized by a Vif from the SIV infecting moustached guenon monkeys (SIVmus), showing that these mutants are still susceptible to Vif proteins adapted towards these interfaces (**Figure 1B**).

Although HIV-1 Vif is sensitive to combinations of mutations at 128 and 130 found in other primates (**Figure 1**), because the natural variation at these sites is limited, these analyses do not address the overall parameters that govern the ability of HIV-1 Vif to recognize human A3G. In order to more globally access the evolutionary pathways available for human A3G to resist antagonism by Vif, we constructed a library of plasmids using degenerate NNS primers to introduce all 20 amino acids to A3G sites 128 and 130 in combination (**Figure 2A**). We screened 244 clones, of which 175 were unique, to test for antiviral activity using a modified single-cycle infectivity assay (**Figure 2B**). In this assay, each A3G mutant was co-expressed in 293T cells with a luciferase reporter HIV-1 molecular clone in which *vif* was deleted or intact. Viruses produced from these cells were then used to infect SupT1 cells to measure the antiviral activities of each A3G mutant in the presence or absence of Vif. Each iteration of the screen contained a vector control to normalize infection values, and a wild-type A3G control to establish the standard deviation of wild-type activity and Vif antagonism. Data generated from this screen are provided in **Supplemental Table 1**.

**Figure 2.**
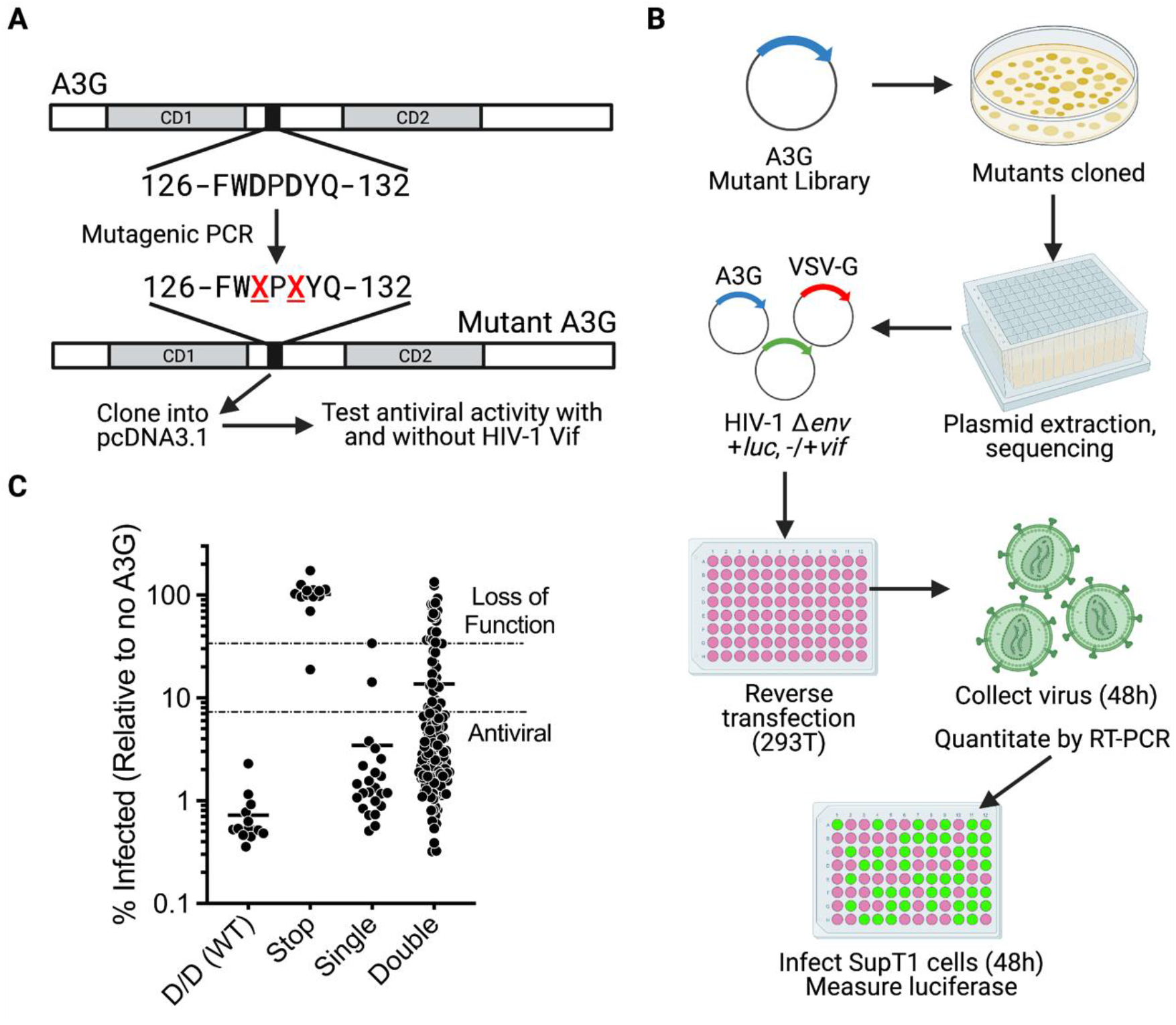
Design and implementation of an A3G combinatorial mutagenesis screen. **A)** A3G editing strategy to introduce mutations at codons 128 and 130. Mutations at codons 128 and 130 were introduced via saturation mutagenesis with primers encoding NNS sequences in place of wild-type A3G codons 128 and 130 (N = any nucleotide, S = Guanine or Cytosine only). Mutant A3G clones were inserted into a transient mammalian expression vector (pcDNA3.1) for downstream applications. **B)** Workflow of A3G combinatorial mutagenesis screen. The A3G mutant library (described in **A**) was transformed into bacteria, plated for single colonies to maintain clone diversity, and grown and extracted via small-scale plasmid preparation. Plasmid concentrations were normalized prior to transfection into 293T cells co-expressing a luciferase-expressing HIV provirus (with and without Vif), and the VSV-G envelope. Virus was propagated for 48 hours prior to collection. Simultaneous infection of SupT1 cells was performed with a standard volume of virus while determining concentration of virus particles via RT-PCR. Extent of infection was measured after 48 hours through luciferase activity, and raw luciferase values were normalized by the RT-PCR-quantified concentration of each mutant A3G-packaged virus. **C)** Single-cycle infectivity assays performed with *vif*-deficient HIV-1 produced in the presence of wild-type A3G (D/D WT), A3G containing premature stop codons at 128 or 130 (Stop), or any variants with single or double mutations screened in this study (Single and Double, respectively). ‘Loss of function’ mutants were defined by having antiviral activity within two standard deviations of A3G containing premature stop codons (top dotted line), and ‘antiviral’ mutants were defined as falling within one log of wild-type A3G activity (bottom dotted line). Solid lines indicate the mean of each group.

In the absence of Vif, wild-type A3G reduced infection to less than 1% relative to vector control (mean 0.724%), while mutants display a wide range of antiviral activity (**Figure 2C, Supplemental Table 2**). Antiviral mutants were defined as those that inhibit *vif*-deficient HIV-1 infection to within one log of average wild-type A3G activity (i.e., mutants that reduce infection to 7.24% or less). As an internal control, mutations containing a premature stop codon lost all antiviral activity, and cells were similarly infected relative to vector control (**Figure 2C**). Loss of function mutants were defined as those that only restrict infection within two standard deviations of the mean activity of stop codon-containing mutants. Of note, most mutants with a single change at 128 or 130 remain antiviral (91%, 21/23 mutants), whereas nearly a third of double mutants (30.3%, 46/152 mutants) fall outside of one log of wild-type activity (**Figure 2C**). This suggests that negative epistasis can exist between these two sites and lower antiviral efficacy.

To understand what mutations may be contributing to reduced A3G activity, we generated difference logo plots to compare sequences in the “loss of function” group to the library of mutants tested, observing an enrichment of proline at 128 and aromatic residues at 128 and 130 (**Figure 3A, Supplemental Table 3**). In agreement with this, all mutants containing D128P lost activity comparable to stop-codon mutants, whereas the same effect is not observed with D130P-containing mutants (**Figure 3B, Supplemental Table 2**). On average, aromatic-containing mutants lose antiviral potency, but with a high degree of variance with no site-specific dependency (**Figure 3B, Supplemental Table 2**). We next tested the effects of D128P, D130P, or aromatic-containing mutants D128Y, D130Y, and D128Y/D130Y to confirm our findings from the screen data analyses. Indeed, the D128P mutant loses all activity compared to wild-type A3G, while the activity of D130P remains at wild-type levels (**Figure 3C**). When introduced singly, tyrosine has no detrimental effect, whereas the double D128Y/D130Y mutant results in a 17-fold reduction in antiviral activity (**Figure 3C**). In all cases, these mutants are susceptible to HIV-1 Vif antagonism, suggesting that these mutations retain some antiviral activity that can be inhibited by HIV-1 Vif. Compared to wild-type A3G, all mutants tested had less expression, with the lowest expression corresponding with poor antiviral efficacy (**Figure 3C**). These results suggest that poisonous mutations and negative epistasis decrease the antiviral efficacy of human A3G through a decrease in expression of the protein, thereby limiting viable paths towards HIV-1 Vif resistance.

**Figure 3.**
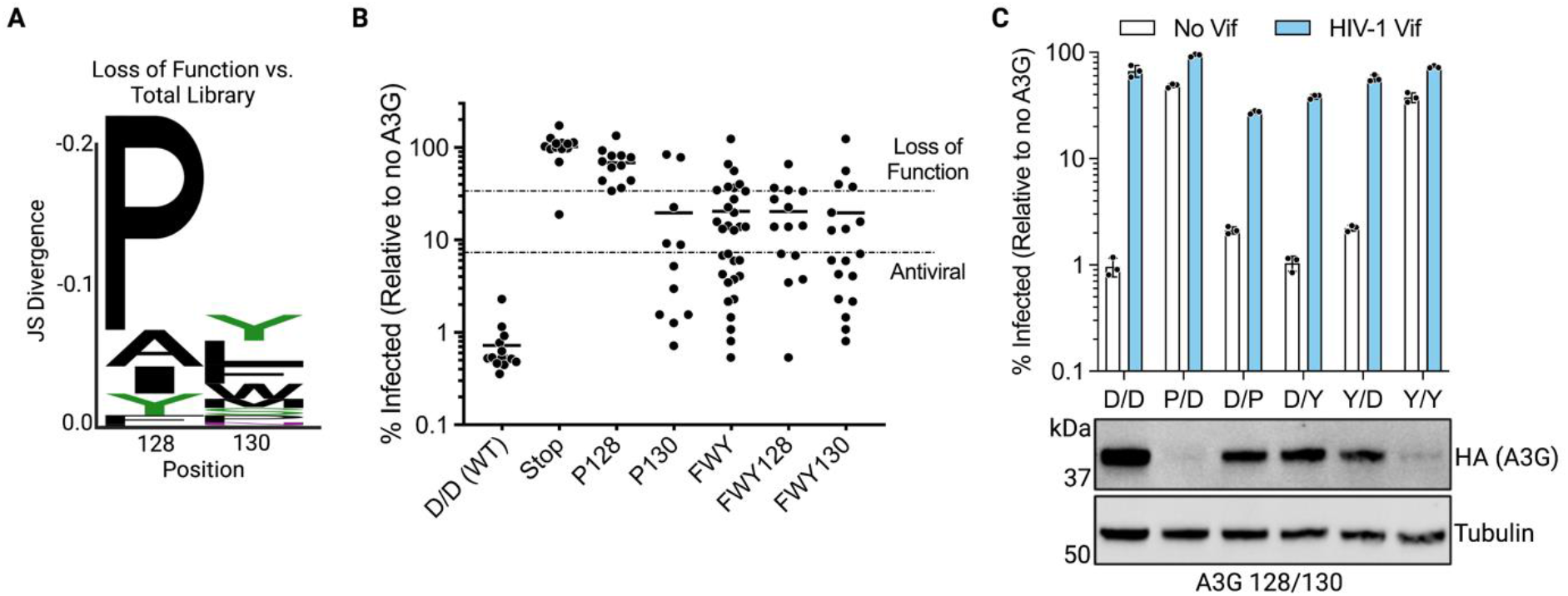
Negative epistasis and “poison” residues in A3G limits paths of Vif resistance. **A**) Difference logo plot representing the enrichment of residues at A3G sites 128 and 130 in mutants identified as ‘loss of function’ compared to the total library of mutants tested. Stack height was calculated using Jensen-Shannon Divergence comparing the amino acid frequencies in each group. Total stack height represents difference between these groups, and individual amino acid heights represent contribution towards the total difference. **B**). Single-cycle infectivity assays performed with *vif*-deficient HIV-1 produced in the presence of wild-type A3G (D/D WT), A3G containing premature stop codons at 128 or 130 (Stop), or mutants containing the indicated motifs. P = proline, FWY = all mutants containing aromatic residues phenylalanine (F), tryptophan (W), and tyrosine (Y), FWY128 = aromatic residues at 128, FWY130 = aromatic residues at 130. ‘Loss of function’ mutants were defined by having antiviral activity within two standard deviations of A3G containing premature stop codons (top dotted line), and ‘antiviral’ mutants were defined as falling within one log of wild-type A3G activity (bottom dotted line). Solid lines indicate the mean of each group. **C**) Top: Validation of selected ‘loss of function’ mutants through single-cycle infectivity assays performed with *vif*-deficient (white) or *vif*-intact (blue) HIV-1 produced in the presence of the indicated A3G mutants. D/D denotes wild-type A3G at positions 128/130. Bottom: Immunoblot of wild-type A3G and ‘loss of function’ A3G mutants. Immunoblotting was performed using anti-HA to detect HA-tagged A3G, and anti-Tubulin as a loading control.

### 2.2. Possible paths of human A3G towards HIV-1 Vif resistance

Mutants in the screen were evaluated both in the absence and presence of Vif in parallel to determine which variants in human A3G could be antagonized by HIV-1 Vif. We evaluated A3G susceptibility towards HIV-1 Vif by comparing the ratio of the percent infectivity in the presence of HIV-1 Vif over the percent infectivity in its absence (termed ‘Fold Vif Antagonism’). We limited our analyses to mutants that restrict infection within one log of wild-type A3G (**Figure 2C**, ‘Antiviral’ mutants) so that only the more antiviral versions of A3G were included (**Figure 4A, Supplemental Table 2**). A3G mutants that were antagonized by HIV-1 Vif to within two standard deviations of the amount of antagonism of wild-type A3G were called ‘Susceptible’ to HIV-1 Vif (**Figure 4A** above the top dotted line), while A3G mutants that were within two standard deviations of the known Vif-resistant mutant D128K were called ‘Resistant’ (**Figure 4A** below the bottom dotted line).

**Figure 4.**
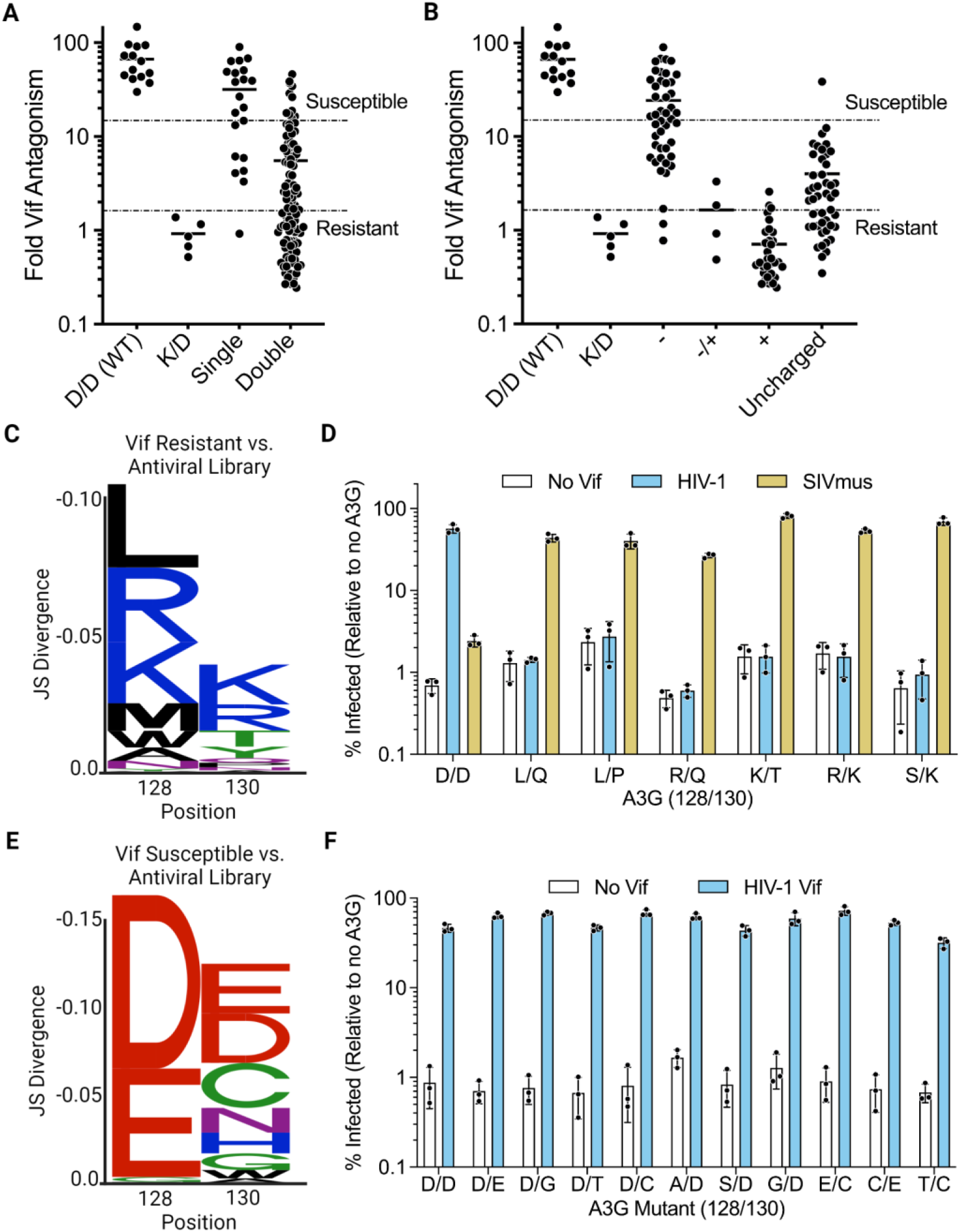
Combinatorial mutagenesis of A3G at 128 and 130 reveals HIV-1 Vif interface flexibility. **A**) Fold Vif antagonism (the ratio of percent infection in the presence of Vif over percent infection in the absence of Vif) was calculated from single-cycle infectivity assays performed with HIV-1 produced in the presence of wild-type A3G (D/D WT), a known Vif-resistant A3G mutant (K/D), and antivirally active screen mutants containing a mutation at either 128 or 130 (Single) or at both sites (Double). ‘Susceptible’ mutants were defined as having a Fold Vif Antagonism within two standard deviations of wild-type A3G (top dotted line), and ‘Resistant’ mutants were defined as falling within two standard deviations of A3G-K/D activity (bottom dotted line). Solid lines indicate the mean of each group. **B**) Fold Vif antagonism from single-cycle infectivity assays performed with HIV-1 produced in the presence of negatively charged 128/130 mutants (-), net neutral mutants containing a positive and a negative charge (-/+), positively charged mutants (+), and uncharged mutants. Solid lines indicate the mean of each group. **C**) Difference logo plot representing the enrichment of residues at A3G sites 128 and 130 in mutants identified as ‘Vif Resistant’ compared to all antivirally active mutants. Stack height was calculated using Jensen-Shannon Divergence comparing the amino acid frequencies in each group. Total stack height represents difference between these groups, and individual amino acid heights represent contribution towards the total difference. **D**) Validation of selected ‘Vif Resistant’ mutants through single-cycle infectivity assays performed with HIV-1 without Vif (white), with Vif intact (blue), or with SIVmus Vif (yellow) produced in the presence of indicated A3G 128/130 mutants. D/D (WT) denotes wild-type A3G. Percent infectivity calculated relative to a no A3G control. Error bars represent the standard deviation of the mean of three biological replicates. **E**). Difference logo plots as in **C** comparing mutants identified as ‘Vif Susceptible’ to all antivirally active mutants. **F**) Validation of selected ‘Vif Susceptible’ mutants through single-cycle infectivity assays performed with HIV-1 without Vif (white) or with Vif intact (blue), produced in the presence of indicated A3G 128/130 mutants. D/D (WT) denotes wild-type A3G. Percent infectivity calculated relative to a no A3G control. Error bars represent the standard deviation of the mean of three biological replicates.

Previous reports indicate that negatively charged residues at A3G 128 and 130 are required for HIV-1 Vif antagonism, while mutation to a single positively charged residue confers resistance [19–21]. However, we also observed that the majority of A3G mutants that are resistant to HIV-1 Vif have mutations at both 128 and 130 rather than mutation at only one of these positions (Figure 4A: compare ‘Single’ versus ‘Double’). Nonetheless, when mutants are segregated by their charge; a net negative charge at both sites results in HIV-1 Vif susceptibility, whereas a net neutral or positive charge confers potent resistance (**Figure 4B, Supplemental Table 2**). However, we also observe uncharged A3G mutants conferring both susceptible and resistance phenotypes (**Figure 4B**). We generated a difference logo plot to compare sequences enriched in mutants that completely resist Vif to all antivirally-active mutants tested, finding a strong preference for positively charged lysine (K) and arginine (R) (**Figure 4C, Supplemental Table 3**). In addition to charged amino acids, we also observed a preference for uncharged residues, such as leucine (L), suggesting that uncharged residues can maintain antiviral potency while simultaneously resisting HIV-1 Vif antagonism.

We validated six Vif-resistant mutants from the screen in single-cycle infectivity assays (**Figure 4D**). We considered four charged mutants, D128R/D130Q (R/Q), D128K/D130T (K/T), D128R/D130K (R/K), and D128S/D130K (S/K), as well as two uncharged mutants D128L/D130Q (L/Q) and D128L/D130P (L/P). These mutants were all capable of maintaining similar antiviral activity both in the absence and presence of HIV-1 Vif (**Figure 4D**), indicating that multiple novel mutations can maintain antiviral efficacy while resisting Vif antagonism. Furthermore, our results with the uncharged L/Q and L/P mutants indicate that charge is not required to resist Vif antagonism. These mutants were still completely susceptible to Vif from the SIV infecting mustached guenons, SIVmus, confirming that these mutants specifically resist HIV-1 Vif (**Figure 4D**).

### 2.3. HIV-1 Vif displays broadened interface specificity towards human A3G mutants

Despite the presence of amino acid changes that confer resistance of human A3G to HIV-1 Vif (**Figure 4C,D**), we find that HIV-1 Vif has considerable flexibility in antagonizing human A3G with different amino acids at positions 128 and 130. A logo plot of those A3G mutants that are antagonized by HIV-1 Vif shows an enrichment in A3G mutants that contain a wild-type aspartic acid (D) at 128 or 130, with the acidic glutamic acid (E) also highly enriched (**Figure 4E, Supplemental Table 3**), consistent with the results in Figure 1. We also observe greater variation and less preference for novel residues at 130, suggesting that HIV-1 Vif can tolerate more changes at A3G 130 than at 128 (**Figure 4E**), and can still antagonize A3G mutants with an uncharged amino acid at position 130.

We validated HIV-1 Vif antagonism towards the most susceptible mutants in our screen, employing mutants containing residues enriched in our logo plot analysis (**Figure 4E, 4F**) including single mutations at 128 (D128A, D128S, D128G) and 130 (D130E, D130G, D130T, D130C), double mutations that maintain negative charge (D128E/D130C, D128C/D130E), and an entirely uncharged mutant (D128T/D130C). We found that all of these mutants were strongly antagonized by HIV-1 Vif, supporting the idea that HIV-1 Vif can tolerate novel mutations at both 128 and/or 130, as well as accommodate completely uncharged mutants (**Figure 4F**). These results suggest that HIV-1 Vif has evolved considerable flexibility in its recognition of human A3G. Importantly, this flexibility contrasts with SIV Vif proteins from Old World monkeys, such as SIVagm.tan Vif, which have strict requirements for both K128 and D130 in their host A3G [15,22].

We further considered the evolutionary pathways by which human A3G remains susceptible or resistant to HIV-1 Vif by considering the effects of possible single-nucleotide mutations at either 128 or 130 (**Table 1**). Although Vif resistance can be achieved, it is mainly accessed through sampling of positively charged lysine or arginine or by mutating away from negatively charged, wild-type aspartic acid residues (**Figure 4B, 4C**), neither codons 128 nor 130 in human A3G can readily access positively charged residues, suggesting that HIV-1 Vif resistance is unlikely to be achieved via single nucleotide polymorphism at either site. This contrasts with the consensus sequence of codon 128 in Old World monkey A3G, which can access a charge-swap mutation, K128E, within a single nucleotide change (**Table 1**). Finally, while residue 130 is conserved between human and Old World monkey A3G, accessible residues do not confer resistance in human A3G due to the broadened specificity of HIV-1 Vif (**Table 1**, **Figure 4E, 4F**). Thus, while it is possible for human A3G to evolve away from HIV-1 Vif, it is evolutionarily difficult to do due so because of access to charge-changing amino acids through single nucleotide mutations and the broad specificity of HIV-1 Vif to additional amino acids at positions 128 and 130.

**Table 1.**
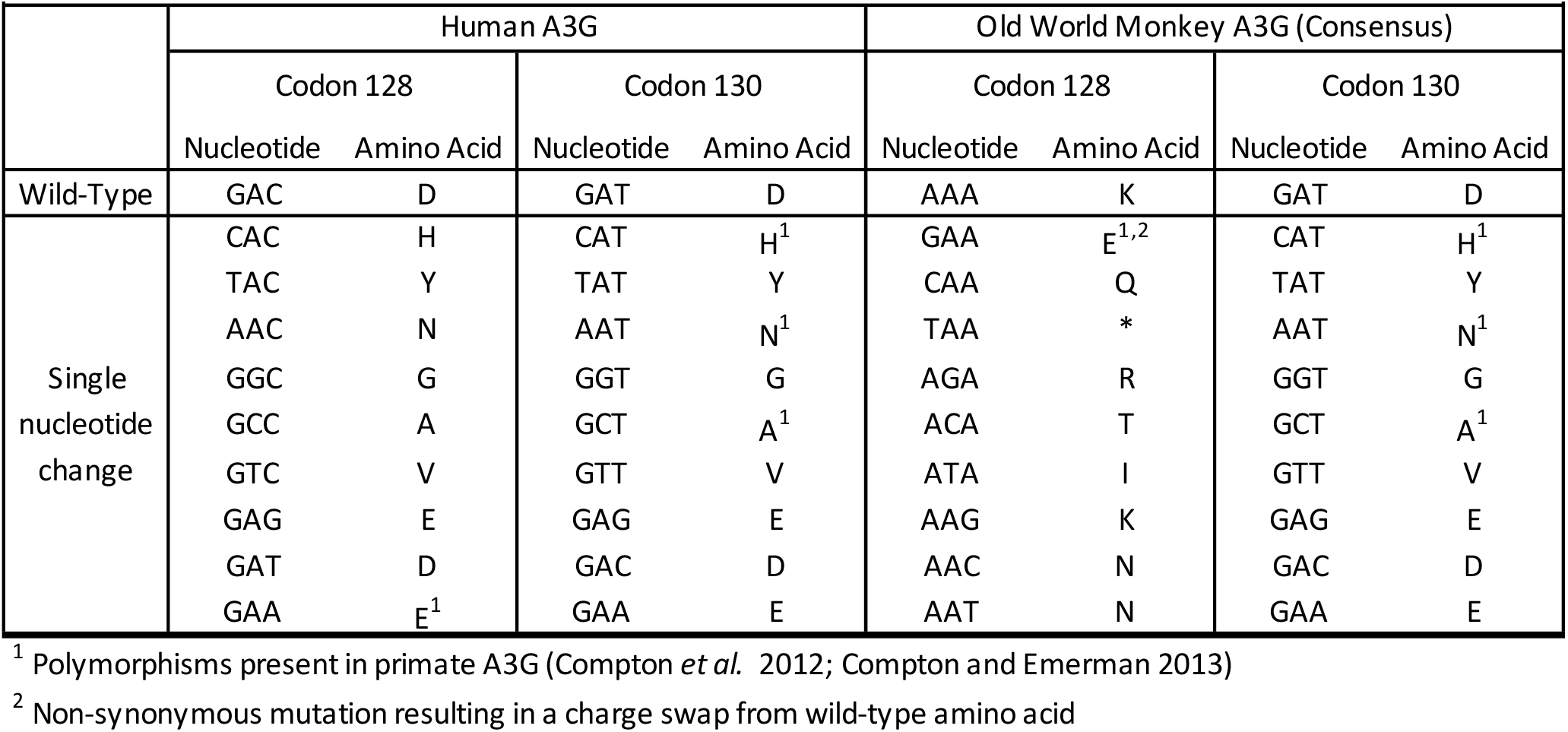
Evolutionary access of amino acids at A3G sites 128 and 130 Analysis of amino acid accessibility via single nucleotide polymorphism of wild-type codons at positions 128 and 130. Human A3G and the consensus sequence of Old World Monkey A3G are compared.

| **Table 1** Evolutionary access of amino acids at A3G sites 128 and 130 |  |  |  |  |  |  |  |  |
| --- | --- | --- | --- | --- | --- | --- | --- | --- |
|  | Human A3G |  |  |  | Old World Monkey A3G (Consensus) |  |  |  |
|  | Codon 128 |  | Codon 130 |  | Codon 128 |  | Codon 130 |  |
|  | Nucleotide | Amino Acid | Nucleotide | Amino Acid | Nucleotide | Amino Acid | Nucleotide | Amino Acid |
| Wild-Type | GAC | D | GAT | D | AAA | K | GAT | D |
| Single nucleotide change | CAC | H | CAT | H<sup>1</sup> | GAA | E<sup>1,2</sup> | CAT | H<sup>1</sup> |
|  | TAC | Y | TAT | Y | CAA | Q | TAT | Y |
|  | AAC | N | AAT | N<sup>1</sup> | TAA | * | AAT | N<sup>1</sup> |
|  | GGC | G | GGT | G | AGA | R | GGT | G |
|  | GCC | A | GCT | A<sup>1</sup> | ACA | T | GCT | A<sup>1</sup> |
|  | GTC | V | GTT | V | ATA | I | GTT | V |
|  | GAG | E | GAG | E | AAG | K | GAG | E |
|  | GAT | D | GAC | D | AAC | N | GAC | D |
|  | GAA | E<sup>1</sup> | GAA | E | AAT | N | GAA | E |

<sup>1</sup> Polymorphisms present in primate A3G (Compton *et al.* 2012; Compton and Emerman 2013)
<sup>2</sup> Non-synonymous mutation resulting in a charge swap from wild-type amino acid

### 2.4. Adaptation of HIV-1 Vif to human A3G broadened its target tolerance

We next asked whether broadened specificity of Vif towards human A3G 128/130 variants occurred during cross-species adaptation to chimpanzee, or from further adaptation from chimpanzee to human. A critical adaptation event for lentiviral Vif towards hominid A3G recognition required a Y86H mutation within loop 5 of SIVrcm Vif to accommodate the D128 residue in chimpanzee A3G ([5], **Figure 5A**). We previously observed that this mutation could fully antagonize both chimpanzee and human A3G while maintaining recognition towards the K128 residue of rcmA3G, suggesting that the Y86H mutation broadened lentiviral Vif specificity [5]. To determine if this same change also allowed the broad specificity of HIV-1 Vif that we observed (**Figure 4F**), we utilized a selection of validated mutants potently antagonized by HIV-1 Vif (E/D, D/G. G/D, E/C, C/E, and T/C) and assessed their antiviral activity in the presence of HIV-1 Vif evolutionary precursors: SIVrcm Vif, and the full loop 5 swap between SIVrcm and SIVcpz Vifs (SIVrcm L5swap) (**Figure 5A, 5B**).

**Figure 5.**
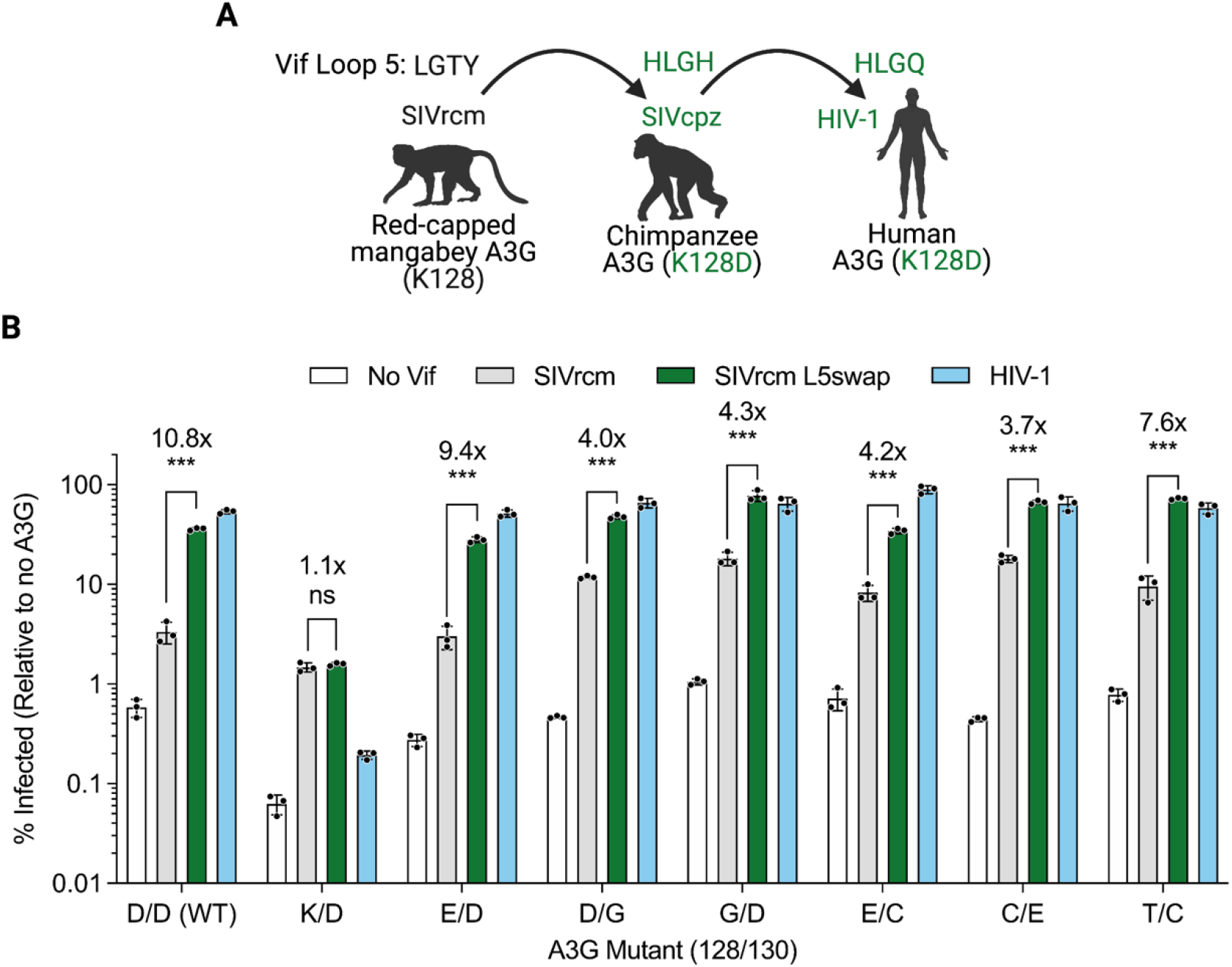
Adaptations in Loop 5 of Vif determine flexibility towards A3G variants. **A**) Schematic outlining the evolutionary trajectory of VIf Loop 5 facilitating the adaptation of lentiviral Vif from Old World monkey to hominid primate A3G, as described previously (Binning *et al*., 2019). **B**) Single-cycle infectivity assays of HIV-1 with *vif*-deficient (white), SIVrcm Vif (gray), SIVrcm Vif 83-LGTY-86 to 83-HLGH-86 (“L5swap,” green), and HIV-1 Vif (blue), produced in the presence of the indicated A3G mutants. Error bars depict the standard deviation of the mean of three biological replicates. Brackets denote the fold difference between infection in the presence of SIVrcm Vif L5swap versus infection in the presence of SIVrcm Vif. *** = p <0.001, ns = not significant, unpaired t test.

In all cases, SIVrcm Vif can weakly antagonize human A3G mutants, regardless of the residues at 128 and 130, due to a conserved hydrophobic anchoring point upstream of Loop 5 [5]. As we found previously, the adaptive SIVrcm L5swap mutant gains strong and specific recognition of the D128/D130 motif found in wild-type human A3G, but does not confer the same adaptation towards the D128K mutant, indicating that SIVrcm Vif Loop 5 accommodates sequence-specific interfaces in human A3G ([5] and **Figure 5B**). Although we notably observe differential potency of wild-type SIVrcm Vif against human A3G mutants, substituting the HIV-1 Vif Loop 5 sequence significantly increases SIVrcm Vif antagonism in each case (**Figure 5B**). Thus, the four amino acids in Loop 5 of Vif are primarily responsible for accommodating interface changes in human A3G 128 and 130. Taken together, our results suggest that the Loop 5 region of Vif may determine its breadth and specificity towards species-specific A3G interfaces, and the broadened specificity observed in HIV-1 Vif was originally conferred in the adaptation of SIVrcm Vif towards chimpanzee A3G. Moreover, the specificity of Vif after cross-species transmission of SIVrcm to chimpanzees broadened beyond just the amino acids naturally found in A3G of extant primate species.

## 3. Discussion

We used a prospective evolutionary analysis of the amino acids in human A3G that are under strong positive selection in primates to probe for resistance strategies against HIV-1 Vif. While there is only a limited range of amino acids that have been sampled by nature at positions 128 and 130, we find that the broadened specificity of Vif towards human A3G developed during the primary adaptation of SIVrcm Vif to chimpanzee A3G, enabling the flexible recognition of evolutionarily accessible residues in hominid A3G. Human A3G could prospectively resist HIV-1 Vif antagonism through mutations not found in extant primates, but it is limited by loss of antiviral activity of some mutations at positions 128 and 130 and by the ability to access resistance mutations through single nucleotide changes. Together, our results highlight the flexibility of Vif adaptation to primate A3G and suggest that adaptation to novel host A3G interfaces can be achieved through broad specificity switches dependent upon adaptation towards a new host A3G interface.

### 3.1. Novel HIV-1 Vif resistance strategies but limited evolutionary access

Our combinatorial mutagenesis approach unveiled many mutants that maintain wild-type levels of antiviral activity while also resisting HIV-1 Vif antagonism, demonstrating that several novel resistance strategies are possible. However, these strategies require introducing a positively charged residue or removing both negative charges through dual mutations at 128 and 130, as HIV-1 Vif is capable of antagonizing mutants maintaining a negative charge at either site (**Figure 4**). Importantly, neither strategy can be achieved through single nucleotide polymorphism (**Table 1**). Thus, breaking HIV-1 Vif antagonism can only be achieved through two or more nucleotide substitutions, significantly raising the barrier to durable resistance. In contrast, the consensus codon sequence of 128 in Old World monkey A3G, AAA (K) can access the K128E charge swap single nucleotide polymorphism, AAA (K) to GAA (E), that is found in nature and potently resists many SIV Vif proteins (**Table 1** and [15,22].

Codons 128 and 130 are conserved in all hominid A3G sequenced to date (Compton and Emerman 2013), suggesting that this evolutionary roadblock to prospective Vif resistance exists in all great ape A3G. The D128 residue is unique to the great apes and arose upon divergence from the lesser apes [15,22]. When only considering sequences found in nature (**Figure 1**), a D128K reversion in human A3G is the most accessible path for HIV-1 Vif resistance but requires an E128 intermediate that remains fully antagonized by HIV-1 Vif (**Table 1**, **Figure 1B, 4E, 4F, 5B**). Therefore, while the D128 residue evades recognition by many Old World SIV Vif proteins, fixation of this residue places hominids at a disadvantage for resisting a fully adapted Vif. Furthermore, previous work demonstrates that HIV-1 Vif can re-establish antagonism of the human A3G K128 escape mutant through adaptation of the 14-DRMR-17 motif [26]. Thus, prospective hominid A3G resistance of Vif is unlikely to be stably achieved through positive selection at sites 128 and 130, placing A3G at a distinct disadvantage in the molecular host-arms race against Vif.

An alternative, evolutionarily accessible Vif resistance strategy may be available to hominid primates through A3G P129. Although conserved in all primates except for gorilla and not under positive selection, previous work demonstrates potent Vif resistance conferred by multiple mutations at this site [17,23,27]. Of these, P129A and P129Q are accessible to human A3G through single nucleotide polymorphisms (CCA to GCA, and CCA to CAA, respectively). When introduced in the human A3G backbone, P129A and P129Q confer resistance towards HIV-1 Vif, demonstrating this resistance strategy is readily accessible for human A3G [17,23]. Gorilla A3G uniquely possesses the P129Q allele that resists antagonism by SIVcpz Vif, and acts as a barrier of SIV cross species transmission between great apes [28]. Recently, it was shown that adaptation of SIVcpz Vif towards gorilla required an M16E mutation in F-box-1 (residues 14 to 17) to overcome the barrier imposed by the P129Q residue in gorilla A3G [28]. This is reminiscent of the single SIVrcm Vif adaptive mutation in Loop 5 (residues 83 to 86 in SIVrcm Vif, 81 to 84 in SIVcpz Vif), Y86H, towards the K128D allele in chimpanzee A3G [5]. Moreover, these regions are adjacent in tertiary structures of both SIVrcm and HIV-1 Vif [5,29], suggesting that these two discontinuous determinants may form a single adaptable interface to accommodate novel A3G substrates.

### 3.2. A3G escape from Vif is limited by poison residues

In addition to the difficulty in achieving resistant human A3G alleles through single point mutations at amino acids 128 and 130, another limitation of Vif resistance in human A3G are poisonous mutations that reduce inherent antiviral potency correlated with a loss of protein expression (**Figure 3**). How mutations at these residues affect protein expression remains unclear. Aside from the possibility that these residues may impact translation, protein folding, or posttranslational stability, these sites also reside in a region of A3G that regulate several aspects important for its activity. Site 128 is part of a putative cytoplasmic retention signal spanning residues 113-128, and certain residues may disrupt this signal and prevent A3G antiviral function [30]. These sites are also directly adjacent to a critical motif, 124-YYFW-127, required for the packaging of A3G into retroviral particles [17]. Finally, this region has also been implicated in nucleic acid binding and A3G dimerization that may be important for its antiviral efficacy [31–33]. Thus, Vif proteins may have evolved to recognize a functionally critical region of A3G that governs its stability, localization, virion packaging, nucleic acid binding, and multimerization, thereby preventing novel escape strategies that would otherwise adversely affect intrinsic A3G function.

### 3.3. Broadened specificity of HIV-1 Vif after cross-species transmission of its precursors

Broadening of Vif specificity was previously observed in primates with polymorphic A3G. African Green Monkey subspecies are polymorphic at A3G sites 128 and 130, and the grivet subspecies expresses either the ancestral K128/D130 motif or the minor allele E128. The resulting Vif of the virus that infects grivet monkeys, SIVagm.gri, is capable of antagonizing all seven haplotypes and all four 128/130 motifs present among the African Green Monkeys, including those not present in the grivet population [15]. The mustached guenon monkey also expresses K128/A130 and E128/A130 motifs [22], and the resulting SIVmus Vif possesses remarkably broad specificity beyond sequences found in nature (**Figure 4D**). Thus, A3G polymorphisms in the host population can train Vif to adopt broad interface specificity.

Here, we demonstrate that broadened Vif specificity can also arise during cross-species transmission. The resolved crystal structure of SIVrcm Vif, and in-depth comparison with HIV-1 Vif, revealed key insights into how cross-species adaptation of Vif proteins can occur. We previously found that both Vif proteins adopt the same overall fold, with key structural differences occurring in loop regions [5]. Of these, the Loop 5 region is primarily responsible for adapting SIVrcm Vif towards antagonism of hominid A3G, with Y86 being the primary amino acid determinant. Intriguingly, the Y86H mutation was able to fully antagonize red-capped mangabey, chimpanzee, and human A3G, suggesting that specificity of SIVrcm Vif was broadened with these mutations to facilitate adaptation to hominid A3G [5]. Indeed, adaptive mutations in Loop 5 of SIVrcm Vif were found to broaden SIVrcm Vif specificity towards novel human A3G interfaces potently antagonized by HIV-1 Vif (**Figure 5**). Thus, the broadened specificity towards human A3G 128/130 mutants observed in HIV-1 Vif was originally conferred during SIVrcm Vif adaptation to chimpanzee A3G, while still maintaining potent antagonism of rcmA3G. It is possible that cross-species adaptation of lentiviruses results in Vif proteins with expanded recognition of A3G interfaces, rather than switching specificity towards a particular interface sequence in the novel host. However, further studies are needed to understand how lentiviral Vif can gain antagonism towards novel host A3G interfaces while still maintaining necessary interactions with other A3 proteins as well as the host E3 ubiquitin ligase machinery. Our work suggests that the broadening of Vif specificity during the cross-species transmission of SIV to hominids was a critical adaptation that not only facilitated potent A3G antagonism, but also prevents prospective Vif resistant mutations from arising at this interface.

## 4. Materials and Methods

### 4.1. Plasmids

N-terminally HA-tagged human A3G was cloned into pcDNA3.1 as previously described [34]. Infectivity experiments were performed using the HIV-1 LAI-based molecular clone pLAIΔenvLuc2Δvif, which has been previously described [35], and L-VSV-G for pseudotyping. SIVrcm Vif and the SIVrcm L5 Swap mutant were ligated into the pLAIΔenvLuc2Δvif provirus using MluI/XbaI restriction sites, as previously described [5]. SIVmus Vif was introduced into the pLAIΔenvLuc2Δvif provirus as previously described [15,22].

### 4.2. Mutagenic library construction and preparation

Mutagenesis and related PCR reactions were performed using Q5 High-Fidelity DNA Polymerase (New England Biolabs, #M0491). The mutant A3G library was constructed via multi-site saturation mutagenesis of codons 128 and 130 using NNS degenerative codons (N = A, T, C, or G, and S = G or C) within the same oligonucleotide (“128-130-NNS-R”: 5’-GCTGCGAAGCGCCTCCTGGTASNNTGGSNNCCAGAAGTAGTAGAGGCGGGC-3’). An initial round of PCR was performed to produce two fragments; the first consisting of the 5’ 450 bp region using the antisense 129-130-NNS-R primer and the flanking 5’ sense primer (“pcDNA-HindIII-HA-A3G-F”: 5’-CTGGCTAGCGTTTAAACTTAAAGCTTGCCACCATGTATC-3’), and the second consisting of the 3’ 767 bp region using an internal sense primer (“A3G-395-F”: 5’-AGGAGGCGCTTCGCAGCCTGTGTCAGAAAAG-3’) and the flanking 3’ antisense primer (“pcDNA-XbaI-A3G-R”: 5’-AGCGGGTTTAAACGGGCCCTTCTAGATTAGTTTTCCTGATTCTG-3’). A second round of PCR was performed with the flanking primers pcDNA-HindIII-HA-A3G-F and pcDNA-XbaI-A3G-R using both fragments as a template to produce a full-length, mutagenized HA-tagged A3G library. Flanking primers contain 20 bp of 5’ and 3’ overlap with the pcDNA3.1 vector for Gibson Assembly (New England Biolabs, #E5510). The resulting library had an estimated complexity of 32^2^ = 1024 codon variants and 20^2^ = 400 amino acid variants. The library was transformed into MAX Efficiency DH5α Competent Cells (ThermoFisher Scientific, #18258012) and plated for single colonies. Single colonies were selected and cultured overnight in Miller’s LB Broth (Corning, #46-050-CM) supplemented with 50 ug/mL carbenicillin, and plasmids were extracted using the QIAprep Spin Miniprep Kit (QIAGEN, #27104). Plasmids were diluted to 40 ng/uL for downstream applications and sequenced by the Fred Hutchinson Cancer Research Center Genomics & Bioinformatics shared resource facility.

### 4.3. Cell lines and transfections

Human embryonic kidney 293T (CRL-3216) and human T-lymphoblast Sup-T1 (CRL-1942) cell lines were obtained from ATCC. Cells were maintained at 37°C and 5% CO_2_ in a humidified incubator. 293T cells were cultured in DMEM, high glucose (Thermo Fisher, 1196592) and SUP-T1 cells were cultured in RPMI-1640 (Thermo Fisher, 11875093). All tissue culture media was supplemented with 10% HyClone Fetal Bovine Serum (GE Healthcare Life Sciences, SH3091003) and 1X Penicillin-Streptomycin (Thermo Fisher, 15140122). Cells were maintained for under thirty passages before returning to a lower-passage stock, and routinely tested for mycoplasma contamination by the Fred Hutchinson Cancer Research Center Specimen Processing/Research Cell Bank core facility.

All transfections were performed using 293T cells and TransIT-LT1 transfection reagent (Mirus, MIR 2305) according to manufacturer’s protocol. For immunoblotting, cells were seeded at 1.5×10^5^ cells/mL in 12-well plates and cultured for 24 h prior to transfection with 1 ug of A3G plasmid and 3 uL transfection reagent per well. For viral infectivity assays, cells were reverse transfected in 96-well plates by complexing 60 ng of pLAIdenvLuc2dVif, 40 ng of A3G, and 10 ng of VSV-G plasmids with 3 uL transfection reagent solution (TransIT-LT1 diluted 10X in serum-free DMEM) for 20 min. Complexes totaled 10 uL and were mixed thoroughly with 3.75×10^4^ cells in 90 uL of complete DMEM. In all cases, transfected cells were further cultured for 48 h prior to harvesting cells or supernatant for downstream applications.

### 4.4. Viral Infectivity Assays

VSV-G-pseudotyped viruses were propagated for 48 h after reverse transfection of 293T cells with pLAIΔenvLuc2 and the indicated A3G or vector control plasmids. Supernatant was collected, transferred to a 96-well V-bottom plate, and spun at 1,000xG for 3 min at room temperature to pellet cell debris. From the clarified supernatant, 5 uL was used to quantitate reverse transcriptase (RT) activity by qPCR [36], and 10 uL was added to freshly-plated SupT1 cells (3.75×10^4^ cells/well) pre-treated with 20 ug/mL DEAE-Dextran. Infected SupT1 cells were cultured for another 48 h and directly lysed with 100 uL of Bright-Glo Luciferase Reagent (Promega, #E2610). Cell lysate was measured for luciferase activity using a LUMIstar Omega microplate luminometer (BMG Labtech). Raw luciferase values were normalized to 2,000 mU RT activity for each virus. Infectivity results from the screen are displayed as the percent infectivity relative to no A3G control, while validated infectivity results are displayed as the mean and standard deviation of three independent experiments.

### 4.5. Immunoblotting

Cells were washed twice with PBS, and pelleted by centrifugation (1,000xG for 3 min at 4°C). Whole cell lysate was extracted on ice with RIPA buffer (50 mM Tris-HCl, pH 7.4, 150 mM NaCl, 1.0% Triton X-100, 0.5% sodium deoxycholate, 1 mM EDTA, and 1 mM MgCl_2_) for 20 min. Lysis buffer was supplemented with cOmplete Protease Inhibitor Cocktail (Roche, 11697498001), 10 uM MG132, and 10 units/mL Benzonase. 10 ug of lysate was resolved on a NuPAGE 4-12% Bis-Tris Protein Gel (Thermo Fisher, NP0336). Immunoblotting was performed using the primary antibodies rabbit anti-HA (Proteintech, 51064-2-AP), and mouse anti-Tubulin (Sigma-Aldrich, #T6199) at a dilution of 1:2000 and 1:5000, respectively. Secondary antibodies StarBright Blue 520 Goat Anti-Rabbit IgG (BIO-RAD, 12005869) and StarBright Blue 700 Goat Anti-Mouse IgG (BIO-RAD, 12005866) were used at a dilution of 1:10,000.

### 4.6. Sequence analysis

Mutant sequences were binned into groups based on antiviral activity in the absence (‘Loss of Function,’ ‘Antiviral,’ and ‘Total’) or presence (‘Vif Susceptible’ and ‘Vif Escape’) of HIV-1 Vif. Amino acid position frequency matrices were calculated for each group of mutants, and used to generate difference logo plots using the DiffLogo R package [37].

### 4.7. Data visualization

All infectivity data were visualized using GraphPad Prism 9. Western blots were prepared using ImageJ software [38]. Figures were arranged using BioRender.com.

## Acknowledgements

We thank members of the labs of Michael Emerman and Harmit Malik for their helpful discussions on this project, and Harmit Malik, John Gross, and Jeannette Tenthorey for critical feedback on this manuscript. We also thank Janet Young and Suzanne Karvonen for technical assistance with this project. NMC was supported by the University of Washington STD/AIDS Research Training Fellowship (NIH/NIAID T32-AI07140), and by a 2021 New Investigator Award from the University of Washington / Fred Hutch Center for AIDS Research (NIH-funded program under award number AI027757). This work was also supported by the HIV Accessory and Regulatory Complexes (HARC) Center (NIH/NIAID P50 AI150476 to ME).

## Supplemental Table Legends

**Supplemental Table 1. Single cycle infectivity results for the A3G 128 and 130 combinatorial mutagenesis screen.** Each mutant screened presented with the mutant identity at residues 128 and 130, % infection in the absence or presence of HIV-1 Vif (‘dVif’ and’ HIV-1 Vif’ columns, respectively). Fold Vif Antagonism, the ratio of % infection with Vif over the % infection without, calculated for each data point. Mutants are sorted in alphabetical order. The % infectivity in the absence of Vif, in the presence of Vif, and Fold Vif Antagonism were averaged in cases were more than one replicate was represented in the screen (‘N’).

**Supplemental Table 2. Source data for scatter plot representations of mutants in screen.** Mutant identities and corresponding % infection and Fold Vif Antagonism data (as applicable) are reported. Mean average and standard deviation calculations for control groups are also shown.

**Supplemental Table 3. Binning of mutants for difference logo analysis.** Mutant identities and corresponding % infection and Fold Vif Antagonism data (as applicable) are reported. Mean average and standard deviation calculations for control groups are also shown. Final list of sequences used for each comparison group in the ‘Difflogo’ sheet.

